# Toxicokinetics for organ-on-chip devices

**DOI:** 10.1101/2024.10.10.617253

**Authors:** Nathaniel G. Hermann, Richard A. Ficek, Dmitry A. Markov, Lisa J. McCawley, M. Shane Hutson

## Abstract

Organ-on-chip (OOC) devices are an emerging New Approach Method in both pharmacology and toxicology. Such devices use heterotypic combinations of human cells in a micro-fabricated device to mimic *in vivo* conditions and better predict organ-specific toxicological responses in humans. One drawback of these devices is that they are typically made from polydimethylsiloxane (PDMS), a polymer known to interact with hydrophobic chemicals. Due to this interaction, the actual dose experienced by cells inside OOC devices can differ strongly from the nominal dose. To account for these effects, we have developed a comprehensive toxicokinetic approach to measure and model chemical-PDMS interactions, including partitioning into and diffusion through PDMS. We use these methods to characterize PDMS interactions for ten chemicals, ranging from fluorescent dyes to persistent organic pollutants to organophosphate pesticides. We further show that these methods return physical interaction parameters that can be used to accurately predict time-dependent doses under continuous-flow conditions, as would be present in an OOC device. These results demonstrate the validity of the methods and model across geometries and flow rates.

## Introduction

Organ-on-chip (OOC) devices are a promising New Approach Method in toxicology. By culturing human cells under microfluidic perfusion, such devices can reproduce human organ responses at a miniaturized scale and thus have advantages in terms of cost, ethical concerns, applicability to human toxicology, and adaptability for high-throughput screening.[1, 2, 3, 4, 5, 6, 7]. Unfortunately, this technology also has a well-recognized drawback: most OOC devices are still made from the elastomer polydimethylsiloxane (PDMS), and PDMS tends to interact with and sequester hydrophobic compounds.[8, 9, 10, 11, 12, 13, 14] When that happens, a compound’s nominal inlet concentration is no longer a reliable measure of its in-device chemical dose. There are three strategies for dealing with this problem: (1) avoid testing of hydrophobic chemicals; (2) mitigate the interactions by modifying PDMS or using a different elastomeric material; or (3) measure and model the interactions to account for them and predict time-dependent *in vitro* concentration profiles throughout such devices. Here, we take the latter toxicokinetic approach. We demonstrate relatively simple methods for effectively measuring chemical-PDMS interactions, which include partitioning at the surface and diffusion into the PDMS bulk.[15, 16] We apply these methods to ten hydrophobic compounds, finding that the partition and diffusion coefficients vary from compound to compound across several orders of magnitude. And finally, we validate these methods and the associated 3D partition-diffusion model by demonstrating its ability to predict thru-device doses under continuous microfluidic perfusion. These methods and models are just initial steps in the larger task of translating *in vitro* dose to equivalent *in vivo* organal dose, i.e., *in vitro*-*in vivo* extrapolation (IVIVE).[17, 7]

We have pursued a toxicokinetic approach because the other two strategies are not always feasible. One may avoid testing hydrophobic chemicals in some cases, but that strategy is unworkable for many applications in environmental toxicology: high hydrophobicity is a key characterisitc of many persistent organic pollutants. For this study, we selected several such compounds: three organophosphate pesticides or pesticide metabolites – chlorpyrifos, paraoxon, and parathion – that have been the subject of prior studies with an organ-on-chip neurovascular unit[18] or microphysiometer[19, 20]; a polycyclic aromatic hydrocarbon, benzo[a]pyrene, previously studied both for its ability to disrupt endocrine signaling in an endometrium-on-a-chip device[21, 22] and as a component of cigarette smoke extract (CSE) studied in a fetal membrane-organ-on-chip[23]; and a pharmaceutical, amodiaquine, that has been studied in a human-airway-on-a-chip.[24, 25]. We complemented this set of toxicants and drugs with four fluorescent dyes – fluorescein, fluorescein-5-isothiocyanate (FITC), rhodamine B, rhodamine 6G – that have been used in several studies as tracers and/or analogs for PDMS interactions.[15, 26, 25] Finally, we included indole, which is fairly water soluble (3.56 mg/mL), relatively non-toxic, and serves as an excellent control that interacts with PDMS quickly, but not prohibitively strongly. As shown in Table 1, each of these ten chemicals has an octanol-water coefficient above the logP = 1.8 threshold where interactions with PDMS become a concern.[13]

**Table 1:**
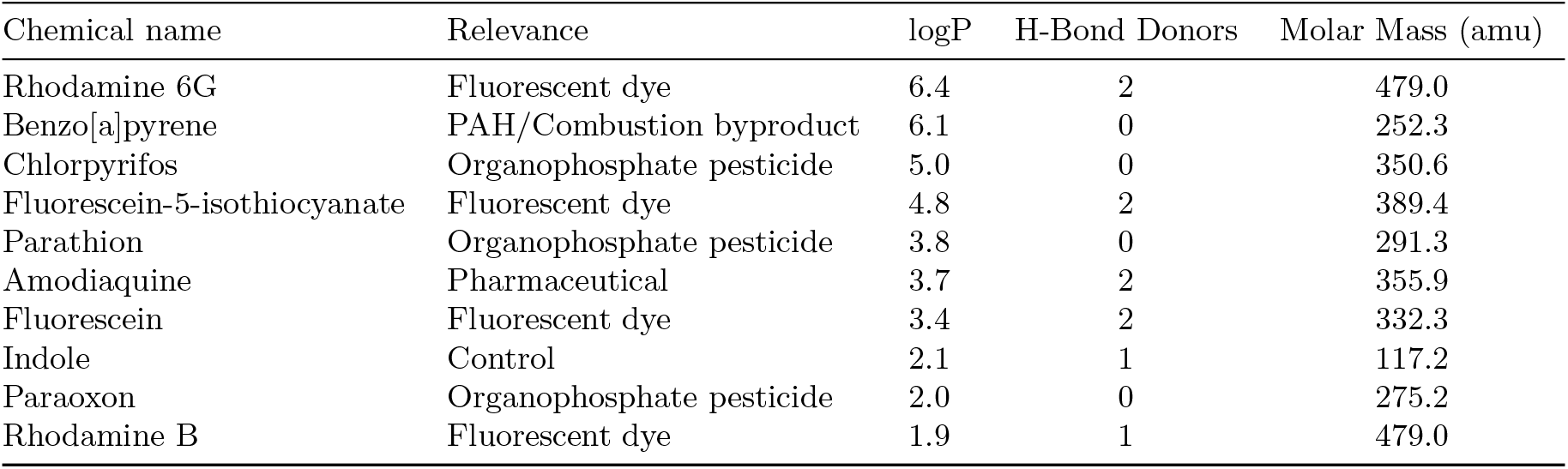
Relevance and selected properties for chemicals tested [34].

As for the strategy of mitigation, several groups are working on alternative materials,[27, 28] but PDMS has many properties that make it ideal for OOC devices including its low cost, flexibility, transparency, gas permeability, and general biocompatibility.[14, 6] It remains the dominant material for microfabricating OOC devices. There are also mitigation techniques that modify the PDMS surface to somewhat limit chemical-PDMS interactions. One popular approach is plasma oxidation, which replaces methyl groups on the polymer chain with hydroxyl groups, making the PDMS surface hydrophilic and eliminating interactions with hydrophobic chemicals. Unfortunately, this change is transient: PDMS will revert over time to a state with an intermediate value of hydrophobicity, which depends on the degree of initial oxidation and subsequent storage conditions.[29, 30, 31] This partial time-dependent reversion is problematic because the interactions with hydrophobic compounds then also change with time. Toxicokinetic modeling would still be needed, and would be more difficult than measuring and accounting for a constant interaction with naive PDMS as we pursue here.

## Methods

### PDMS Preparation

PDMS Sylgard 184 (Dow Corning, Auburn, MI) was mixed in a 10:1 mass ratio of elastomer base to curing agent. To make PDMS disks, PDMS was cast in a 6-mm thick layer, which was then cured overnight in a 67°-C oven. After curing, disks were prepared with a 3-mm radius biopsy punch. These PDMS disks were then annealed for 4 hours in a 200°-C oven to stabilize mechanical properties.[32] PDMS membranes were spun out from small volumes of PDMS to 80-µm thickness and then cured. PDMS with microchannels, measuring 21.1 mm in length, 1.5 mm in width, and 100 µm in height, was made by casting 6 mm of PDMS over a SU-8 photoresist mold. The cast PDMS was then allowed to cure, and annealed. For experiments, each block of PDMS with microchannels was clamped onto a 2 inch by 3 inch microscope slide.

### Chemical Preparation

Chemicals were acquired either as powders or liquids from Sigma Aldrich (St. Louis, MO). Stock solutions were prepared with 1x pH-7.4 phosphate buffered saline (PBS) (Thermo Fisher, Waltham, MA) to near maximum solubility. For chemicals with very low solubility in water, dimethyl sulfoxide (DMSO) was added to increase solubility. DMSO does not partition into nor interact with PDMS, making it a compatible organic solvent[33]. For experiments, stock solutions were diluted in their respective mixed solvent to obtain a peak UV-Vis absorbance close to one.

### Measuring chemical loss to PDMS in disk-soak experiments

Disk-soak experiments were performed by monitoring loss of chemical from solution over time while in contact with a PDMS disk. Concentrations were estimated via UV-Vis absorbance spectra measured with a NanoDrop One^C^ UV-Vis Spectrophotometer (Thermo Fisher, Waltham, MA).

Disk soak experiments were conducted with 2 mL of chemical solution in Type 1P disposable UV plastic cuvettes (FireflySci, Northport, NY). Disks were gently placed on top of this solution such that they floated with only the top surface above the solution (Fig. 1A). Cuvettes were sealed with tight fitting PFTE covers to prevent evaporation. These cuvettes had spectra measured in the spectrophotometer at pseudo-logarithmic times over 48 hours to record concentration loss, while the remainder of the time they were left on a blot mixer to ensure solutions remained well mixed.

**Figure 1:**
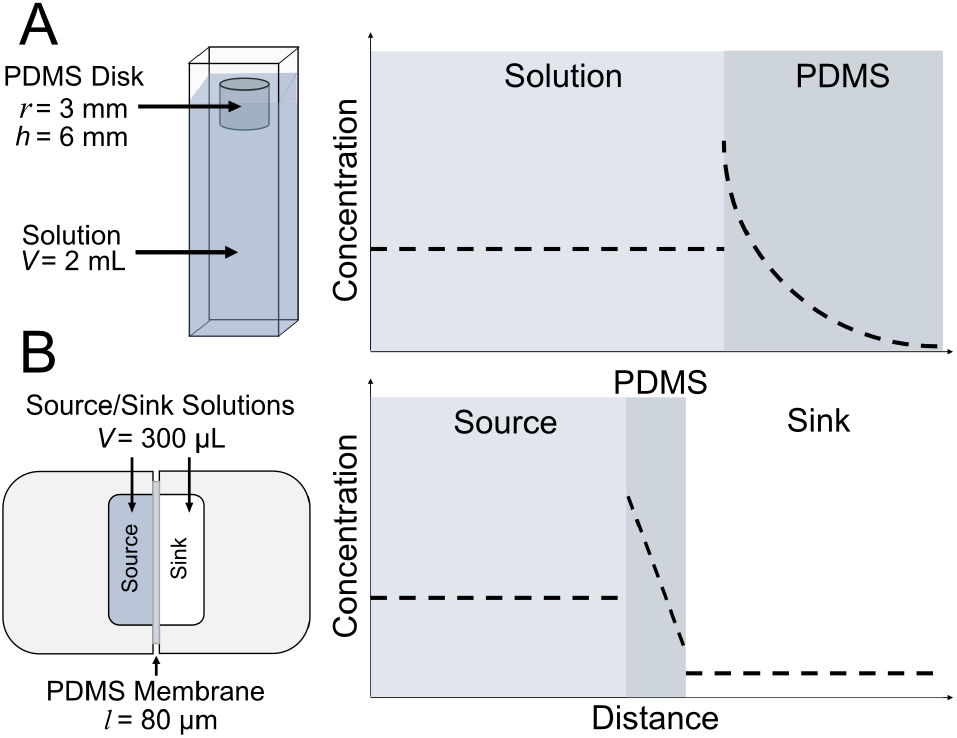
Experimental setups. (A-left) Schematic of a disk-soak experiment in which a disk of PDMS is floated in a cuvette containing a chemical solution of interest; (A-right) An example pre-equilibrium profile of chemical concentration in the solution and PDMS disk. (B-left) Schematic of a membrane experiment in which a thin PDMS membrane separates a source chamber loaded with a chemical solution of interest and a sink chamber loaded with matching solvent; (B-right) An example preequilibrium profile of chemical concentration in source chamber, PDMS membrane, and sink chamber.

Simultaneously, a cuvette of solution with no disk was monitored to control for any chemical interaction with UV plastic. After 48 hours, disks were removed, dried, and placed in cuvettes filled with fresh solvent. These cuvettes were then sealed and monitored for 48 hours to track chemical release from PDMS.

### Measuring chemical diffusion through PDMS

Diffusion-through-membrane experiments were carried out in aluminum devices fabricated in-house. Each device had source and sink reservoirs that were separated by an 80-µm thick PDMS membrane. To prevent leaks, the PDMS-aluminum interface was sealed with high vacuum grease (Dow Corning, Midland, MI). The source and sink were loaded with 200 µL of either chemical solution or matching solvent, respectively. A piece of Scotch tape was used to seal the top of the reservoirs to prevent evaporation (Fig. 1B). For chemicals with reasonable solubility and strong UV/Vis absorbance, concentrations for the source and sink were measured from UV-Vis absorbance spectra of 2-µL aliquots using the attenuated total reflectance pedestal of the NanoDrop One^C^ spectrophotometer. For chemicals with weaker absorbance, the full 200 µL was removed from each chamber, placed in a cuvette for measurement of its UV-Vis absorbance spectrum, and then returned to its original chamber. For most chemicals, measurements were taken at pseudologarithmic intervals over 24 hours. For those chemicals that exhibited fast diffusion, measurements were then repeated at ten minute intervals over 1 hour.

### Direct optical measurement of diffusion in PDMS

For select fluorescent dyes, diffusion in PDMS was also assayed via direct optical visualization. To do so, a 21.1-mm long by 1.5-mm wide by 100-µm tall microchannel in PDMS was filled with a solution of a fluorescent dye, and imaged for three hours using a 1× objective on a Nikon Ti2 Eclipse with X-light V2 spinning disk confocal microscope (Nikon Instruments, Melville, NY). After this time, the microchannel was emptied and dried. The walls of the dry channel were then imaged for an additional 12 hours to follow the diffusive spread of any dye that had previously partitioned into the PDMS. The spatial profile of diffusing dye was fit to a solution to the 1D diffusion equation to estimate diffusivity in PDMS, D_*P*_.

### Measuring chemical loss under constant pumped flow

To replicate chemical loss in a continually perfused PDMS device, we measured chemical loss to PDMS in microchannels perfused at constant flow rates of 5 or 10 µL/min. Perfusion was controlled by a Hamilton Standard Infuse/Withdraw Pump 11 Pico Plus Elite Programmable syringe pump (Harvard Apparatus, Hollison, MA) with 2.5 mL Hamilton syringes (Hamilton, Reno, NV) coupled to the microchannels using Tefzel tubing (McMaster-Carr, Elmhurst, IL). Effluent was collected in 10-min intervals and chemical concentration in each effluent sample was measured via UV-Vis absorbance using the pedestal of the NanoDrop One^C^ spectrometer.

### Simulating experiments and fitting experimental data

Disk-soak, membrane and pumped-flow experiments were simulated using the finite element modeling (FEM) software COMSOL Multiphysics (COMSOL, Inc., Burlington, MA). Disk-soak and pumped-flow experiments were simulated in full three dimensions; simulations of membrane experiments could be reduced by symmetry to just one dimension.

Data from disk-soak and membrane experiments were fit to results from the FEM simulations to estimate chemical-specific model parameters. To make these estimates, simulations of both experiments were run across a uniformly log-spaced grid of all model parameters. This set of 4802 simulations was used to construct a first-order interpolation function, and the experimental concentration data were then fit to this function. Construction of the interpolation function and its regression against experimental data was performed in Mathematica (Wolfram, Champagne, IL).

## Results

We tested ten hydrophobic chemicals (logP *≥* 1.9) with both disk-soak and membrane experiments to measure each chemical’s interactions with PDMS. Seven of these chemicals were sufficiently watersoluble to make measurements directly in phosphate-buffered saline (Fig. 2 and Fig. S1). An additional three were insufficiently water-soluble and required the addition of *≥* 40% DMSO by volume (Fig. 3A). These chemical’s interactions were remarkably varied: some did not partition into PDMS at any detectable level (e.g., FITC in Fig. 2), but others partitioned very strongly, depleting the concentration in solution by up to 80% (e.g., parathion in Fig. 3A); with similarly wide variation, some chemicals diffused through an 80-µm thick PDMS membrane as fast as 1-5 hrs (e.g., indole and paraoxon in Fig. 2), while others needed 12-24 hrs or longer (e.g., amodiaquine and rhodamine B in Fig. 2), and at least one bound to PDMS surfaces, but did not diffuse into the PDMS bulk (rhodamine 6G, Fig. S3). Note that many of the membrane experiments reached final equilibria in which the sum of the source and sink concentrations was less than the initial concentration, revealing that a substantial fraction of the chemical remained in the thin PDMS membrane (e.g., parathion and benzo[a]pyrene in Fig. 3A).

**Figure 2:**
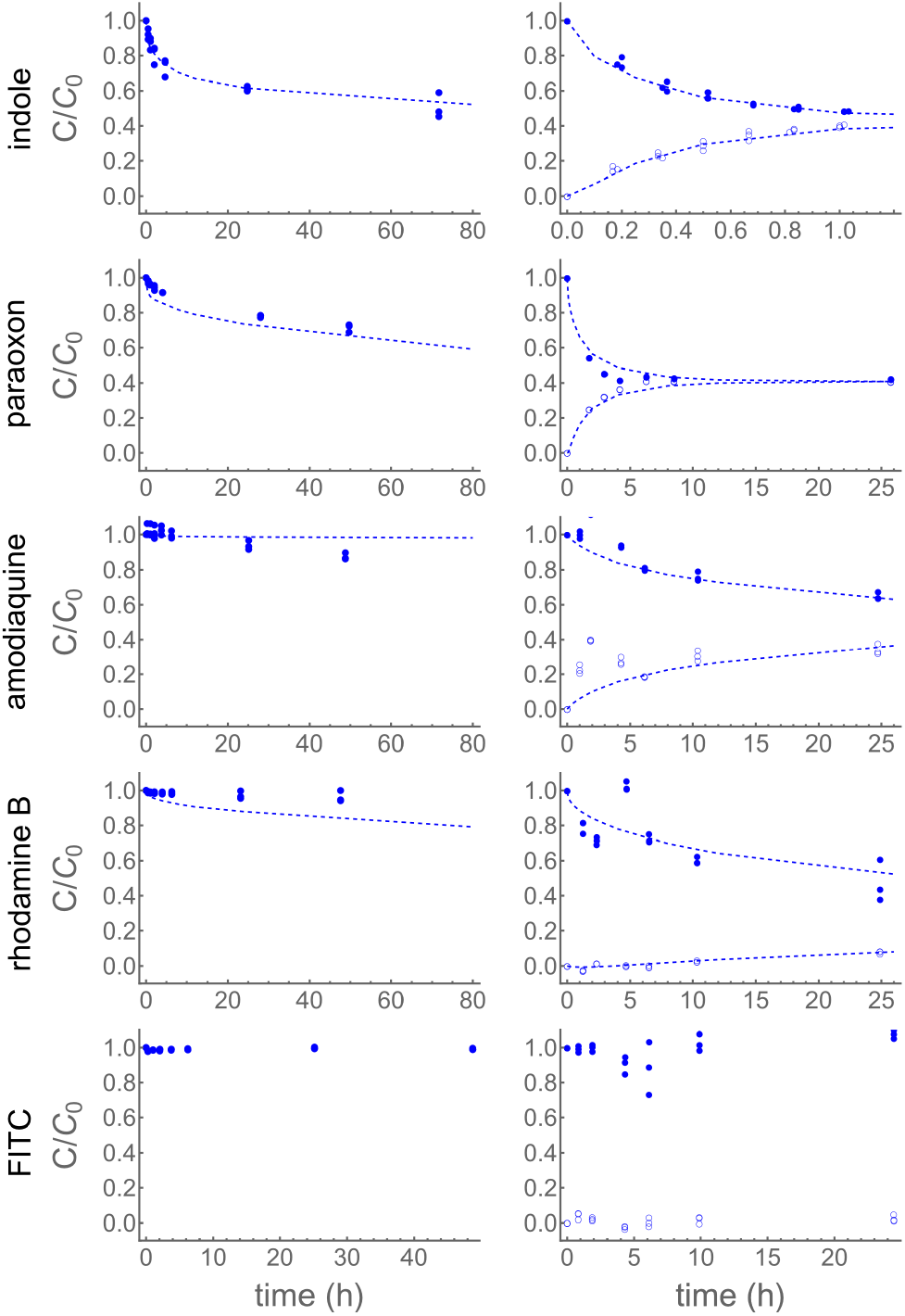
Disk-soak and membrane experiment results for chemicals in solutions of PBS. Left column shows disk-soak data and associated fits (dashed). Right column shows membrane data from both source (filled) and sink (empty) with associated fits (dashed). Since FITC demonstrates no interaction, no fit is shown.

**Figure 3:**
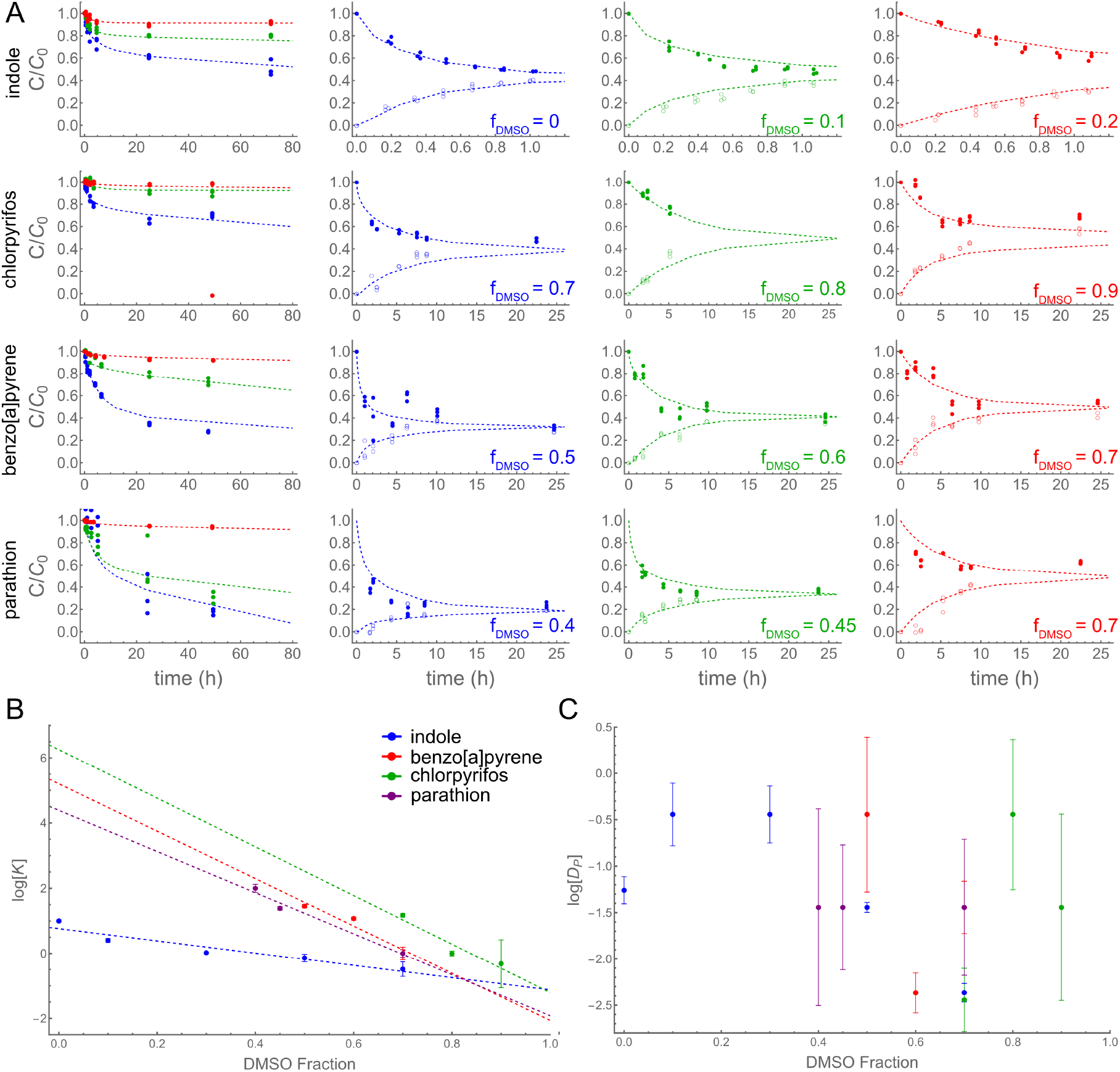
Disk-soak and membrane experiment results for chemicals with added DMSO cosolvent. (A - left column) Disk-soak data for each chemical in mixed solvents at three different DMSO volume fractions, color-coded to match the three right columns with blue, green, and red running from low to high DMSO fraction. (A - right columns) Source and sink data (closed and open symbols) from membrane experiments for each chemical at the noted DMSO volume fraction. Fits shown as dashed lines. (B) Best-fit values of log K versus DMSO volume fraction and linear fits (dashed lines) used to extrapolate to the PDMS-water partition coefficients, log K_*P W*_. (C) Best-fit values of log D_*P*_ versus DMSO volume fraction. Legend in B applies to B and C.

### Parameterizing and modeling chemical-PDMS interactions

To be more broadly useful, measurements of chemical-PDMS interactions must be parameterized with a model that can be extended to other PDMS geometries and solution flow rates. In prior work, we parameterized chemical-PDMS interactions with a surface binding model, which fit disk-soak data well,[13] but with unrealistically large estimates of surface binding capacities. This shortcoming implies diffusion of chemicals into the PDMS bulk and leads the model to make inaccurate predictions when this diffusion becomes important - e.g., long-duration experiments in which a chemical solution continuously flows through microchannels in a large block of PDMS.

We thus sought to parameterize chemical-PDMS interactions with a more accurate and physically realistic model that includes both partitioning at solution-PDMS interfaces and diffusion through bulk PDMS. This model describes the concentration of a chemical species in solution, c_*S*_, and in PDMS, c_*P*_ in any spatial environment with solution-PDMS contact. The concentrations evolve over time as described by the equations

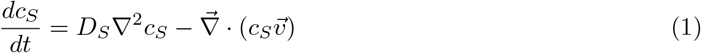

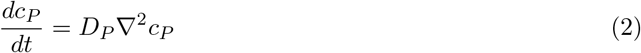

with boundary conditions imposed at the solution-PDMS interface, ∂:

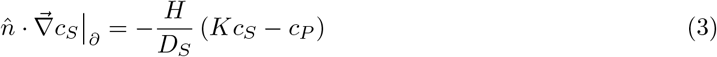

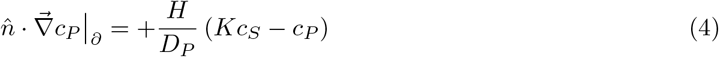

As such, a chemical’s interaction with PDMS is characterized with four parameters: two diffusion constants, D_*S*_ and D_*P*_, which describe diffusion of the chemical in solution and in PDMS respectively; a mass-transfer coefficient at the interface, H; and the solution-PDMS partition coefficient, K. Any flow in the system is described as 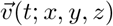, which is zero in the case of both disk-soak and membrane experiments.

In the case of membrane experiments, we extend the model to consider concentrations in both the source and sink chambers. The equations remain similar, but with two solution concentrations, c_*S*_1 and c_*S*_2, representing the source and sink respectively:

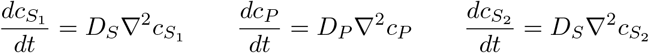

and boundary conditions imposed on both the source-PDMS boundary ∂_1_ and PDMS-sink boundary, ∂_2_:

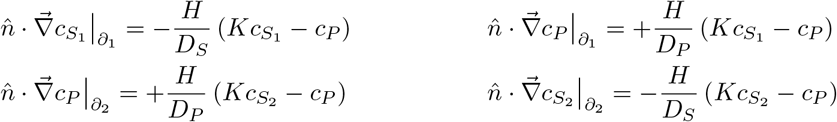

To fit our experimental data to this model, we first ran a set of 4802 numerical simulations for disk-soak and membrane experiment geometries in COMSOL Multiphysics. The simulations were performed over a log-scaled grid of the four parameters: specifically, the parameters spanned the ranges − 2.44 ≤ log D_*S*_ ≤ 3.56, − 6.44 ≤ log D_*P*_ − 0.44, −4.44 ≤ log H ≤ 1.56, and −2 log ≤ K ≤ 4 (D_*S*_ and D_*P*_ in units of mm^2^/h, H in mm/h).The numeric model outputs were then used to construct first-order interpolation functions of the form c_*i*_(D_*S*_, D_*P*_, H, K, t), with concentrations c_*i*_ being those for either disk-soak or membrane experiments. Experimental data were then fit to the interpolation functions to estimate the best-fit parameters for each chemical-solvent combination. We found that the best parameter estimates were provided by fitting both disk-soak and membrane data simultaneously with a shared set of parameters. If either experiment were fit separately, there were near degeneracies in the parameters. For example, fitting disk-soak experiments alone would reasonably constrain the product 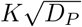, but not the individual parameters K and D_*P*_. Performing simultaneous fits to both experiments relieves the degeneracy.

Of the seven sufficiently water-soluble chemicals tested here, four partitioned into and diffused through PDMS. The data and fits for these four chemicals are shown in Fig. 2, with the best-fit parameters compiled in Table 2. Three of the four (rhodamine B, paraoxon, and indole) partitioned favorably into PDMS (K_*P W*_ *≈* 68, 12 and 8.1, respectively), while amodiaquine partitioned quite weakly (K_*P W*_ *≈* 0.14). A different set of three (indole, paraoxon, and amodiaquine) diffused through PDMS at similar rates with D_*P*_ = 0.01 to 0.06 mm^2^/h, while rhodamine B diffused much slower (4 × 10^*−*4^ mm^2^/h). Note that even the fast-diffusing chemicals have diffusion constants in PDMS that are two orders of magnitude slower than those in water.

**Table 2:** Best-fit parameter values for the partition-diffusion model.

| Chemical | $\log K_{PW}$ | $\log K_{PD}$ | $\log D_P \text{ (mm}^2 \text{ h}^{-1}\text{)}$ | $\log D_S \text{ (mm}^2 \text{ h}^{-1}\text{)}$ | $\log H \text{ (mm h}^{-1}\text{)}$ |
| --- | --- | --- | --- | --- | --- |
| Chlorpyrifos | $6.25 \pm 1.98$ | $-1.21 \pm 0.53$ | $-1.51 \pm 0.30$ | $\geq 3.56$ | $0.04 \pm 0.09$ |
| Benzo[a]pyrene | $5.20 \pm 1.22$ | $-2.06 \pm 0.82$ | $-1.42 \pm 0.30$ | $1.89 \pm 0.26$ | $-0.49 \pm 0.17$ |
| Parathion | $4.39 \pm 0.50$ | $-1.92 \pm 0.47$ | $-1.40 \pm 0.50$ | $\geq 3.56$ | $-1.05 \pm 0.14$ |
| Rhodamine B | $1.83 \pm 0.10$ | – | $-3.44 \pm 0.68$ | $-1.16 \pm 0.12$ | $\geq 1.56$ |
| Paraoxon | $1.08 \pm 0.03$ | – | $-1.96 \pm 0.05$ | $0.52 \pm 0.23$ | $\geq 1.56$ |
| Indole | $0.91 \pm 0.03$ | $-1.12 \pm 0.26$ | $-1.25 \pm 0.09$ | $0.56 \pm 0.07$ | $\geq 1.56$ |
| Amodiaquine | $-0.84 \pm 0.63$ | – | $-1.29 \pm 0.58$ | $\geq 3.56$ | $\geq 1.56$ |

### Modification of chemical interaction by cosolvent

Three of the tested chemicals (benzo[a]pyrene, chlorpyrifos, and parathion) were too poorly soluble to be spectroscopically detectable in pure PBS. We thus added DMSO as a cosolvent. Adding cosolvent affects the partitioning of a chemical between the now mixed solvent and PDMS, but this effect can be accounted for using an extension of the log-linear solubility model of Yalkowsky[35]:

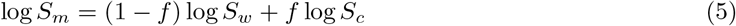

where S_*w*_, S_*c*_, and S_*m*_, are the solubility of a chemical in water, neat cosolvent, and a mixed solution, respectively; and f is the volume fraction of cosolvent in the mixed solution. Since the partition coefficient of a compound between two media, A and B, is related to its solubility in these media,

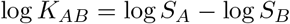

one can derive an equivalent log-linear model to describe the partition coefficient in mixed solvent:

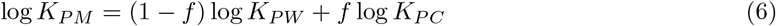

where K_*P W*_, K_*P C*_, and K_*P M*_ are the partition coefficients between PDMS and water, neat cosolvent, or mixed solvent, respectively. log K_*P M*_ should thus be a linear function of the cosolvent fraction. For each of the three chemicals with poor water solubility, we conducted disk-soak and membrane experiments at three different DMSO fractions (Fig. 3A), fit the data in each mixed solvent to the partition-diffusion model, and used the log-linear model to estimate the partition coefficient expected in pure PBS (Fig. 3B). To verify the general validity of the log-linear model, we conducted similar experiments using water-soluble indole at four DMSO volume fractions (Fig. 3A-B). The log-linear extrapolation for indole estimated its log K_*P W*_ as 0.76 *±* 0.14, which agrees with value of 0.91 *±* 0.03 measured directly in pure PBS. With the caveat that the log-linear model likely only provides an order-of-magnitude estimate of log K_*P W*_, we used it to extrapolate the PDMS-water partition coefficient for each of the poorly soluble chemicals (Fig. 3B), and present these values as log K_*P W*_ in Table 2. For each of the poorly water-soluble chemicals, the PDMS-water partition coefficients were estimated to be in the range of 400 to 2000, which is one to two orders of magnitude greater than those of the water-soluble chemicals. The corresponding estimates in pure DMSO, log K_*P D*_ = log K_*P C*_, are also presented in Table 2 to allow interpolation to any DMSO fraction.

To estimate the other model parameters, we simply report the mean of their values obtained at various cosolvent fractions (Table 2). This approach seems very reasonable for diffusivity in PDMS, D_*P*_, which should be independent of any cosolvent present in the solution phase. Consistent with this expectation, estimates of D_*P*_ in different mixed solvents do not show any clear trend with DMSO fraction (Fig. 3C). The average value of D_*P*_ for these chemicals is fairly consistent, around 0.035 mm^2^/h. As for the diffusivity in solution, D_*S*_, one might expect a dependence on cosolvent, but no such dependence is apparent in the estimated values (Fig. S2A). Finally, for the mass-transfer rate, H, the estimates hint at a relationship with DMSO fraction (Fig. S2B), but the data we have collected is too sparse to make any definitive claims and we thus report only the mean values.

### Direct imaging of diffusion in PDMS

To validate the estimates of D_*P*_ obtained from fits to the 3D partition-diffusion model, we directly imaged the diffusion of fluorescent dyes into bulk PDMS. If a dye is distributed with an initial 1D fluorescence profile u(x, 0) that is well approximated by a Gaussian of 1/e^2^-width σ_0_, then its subsequent time-dependent profile under Fickian diffusion with no-flux boundary conditions is given by

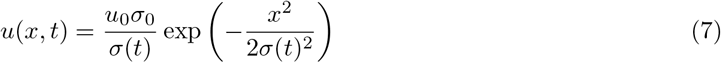

where

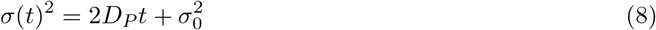

To match these conditions, we loaded microchannels with dye solutions, allowed the dye to diffuse into the channels’ PDMS walls for three hours, emptied the microchannels, and then imaged the diffusive spread of preloaded dye further into the PDMS bulk - without the complicating effects of further partitioning. For each dye, at each time t, we fit the fluorescence intensity profile to equation to estimate the Gaussian square width, σ^2^. We then used a linear fit of σ^2^(t) to equation (8) to estimate D_*P*_. Among the four dyes in our test set, only rhodamine B and 6G bind to or partition into PDMS. We directly imaged the diffusion of both. Rhodamine 6G does not measurably diffuse beyond the PDMS surface, but rhodamine B clearly does, spreading from a width of *∼* 50 µm to *∼* 150 µm over 12 h (Fig. 4AB, Fig. S4). When measured in this direct manner, we estimate the diffusion constant of rhodamine B in PDMS as logD_*P*_ = *−*3.907*±*0.004 (in mm^2^/hr; Fig. 4C), which agrees with the best-fit value found indirectly via disk-soak and membrane experiments, logD_*P*_ = −3.44 *±* 0.68. In similar agreement, the inability of rhodamine 6G to diffuse further into PDMS matches its inability to diffuse through a thin PDMS membrane (Fig. S1).

**Figure 4:**
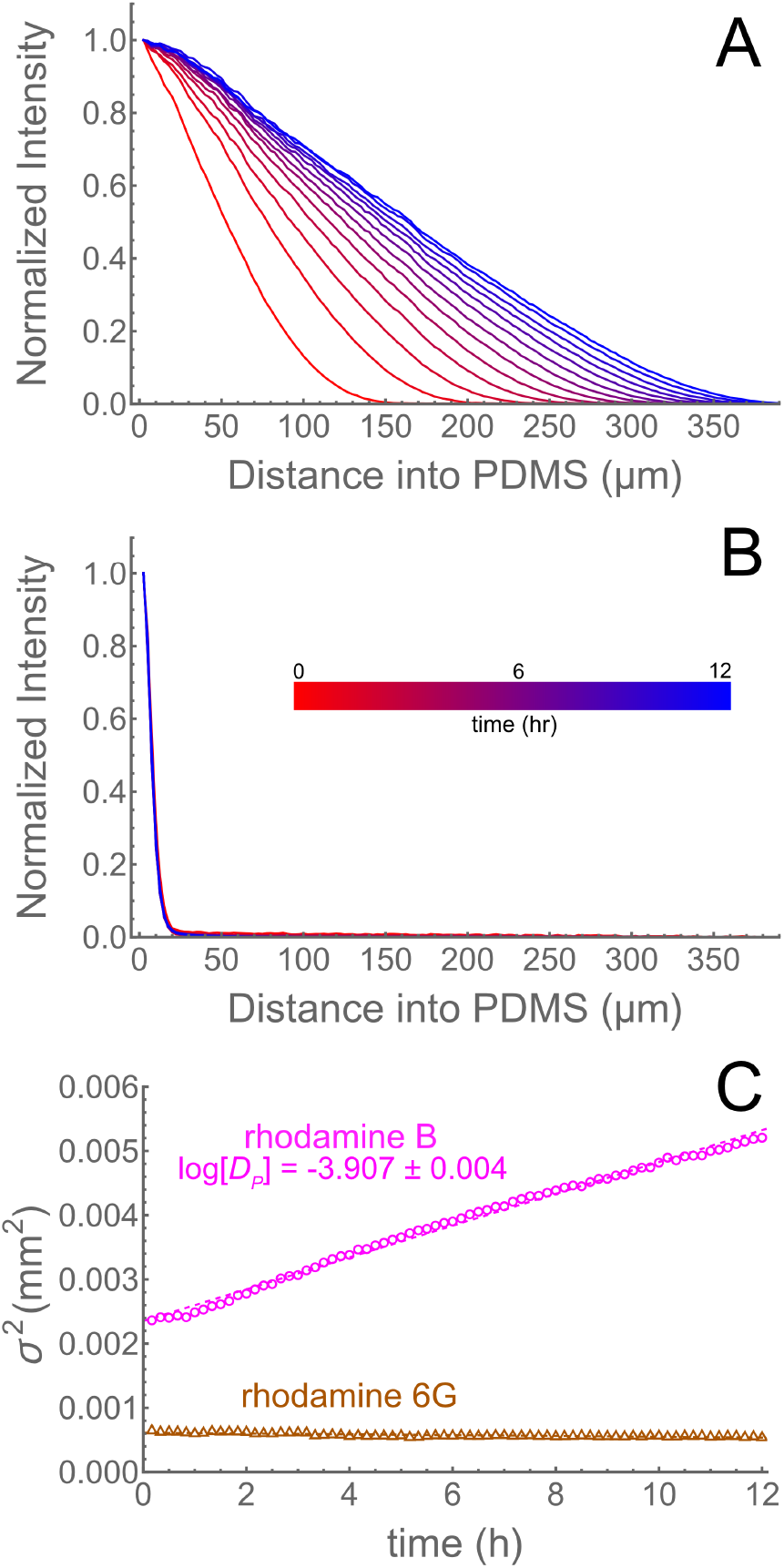
Direct imaging of fluorescent dye diffusion into bulk PDMS. (A) The fluorescence intensity profile of rhodamine B spreads diffusively into PDMS over 12 h. (B) In contrast, the fluorescence intensity profile of rhodamine 6G is essentially static over 12 h, indicating a lack of diffusion into PDMS. (C) For rhodamine B, the Gaussian square width, σ, grows linearly, with linear regression yielding a best-fit value of D_*P*_ (as labeled, in units of mm^2^/h) that agrees with that found indirectly in Table 2. For rhodamine 6G, the Gaussian square width does not measurably change with time.

### Loss of indole during continuous flow in microchannels

To illustrate the applicability of the 3D partition-diffusion model across geometries and flow rates, we used COMSOL Multiphysics to simulate continuous pumped flow of an indole solution through rectangular PDMS channels and compared the results with data from matching experiments. In this case, the parameters for indole were obtained from disk-soak and membrane experiments (Table 2) and used without modification or fitting to directly simulate the first three hours of continuous flow of indole through a microchannel at two flow rates, Q = 5 and 10 µL/min (Fig. S5). In matching experiments, an indole solution was pumped through a microchannel at the same flow rates, and effluent was collected in bins across the duration of the experiment. For the higher flow rate, the simulations slightly underestimate the experimental outlet concentrations in the first hour, but are a good match at longer times; for the lower flow rate, the simulations and experiments are a good match at all times (Fig. 5A). This agreement illustrates the physicality and wide applicability of the model and the methods presented here for inferring its chemical-specific model parameters.

**Figure 5:**
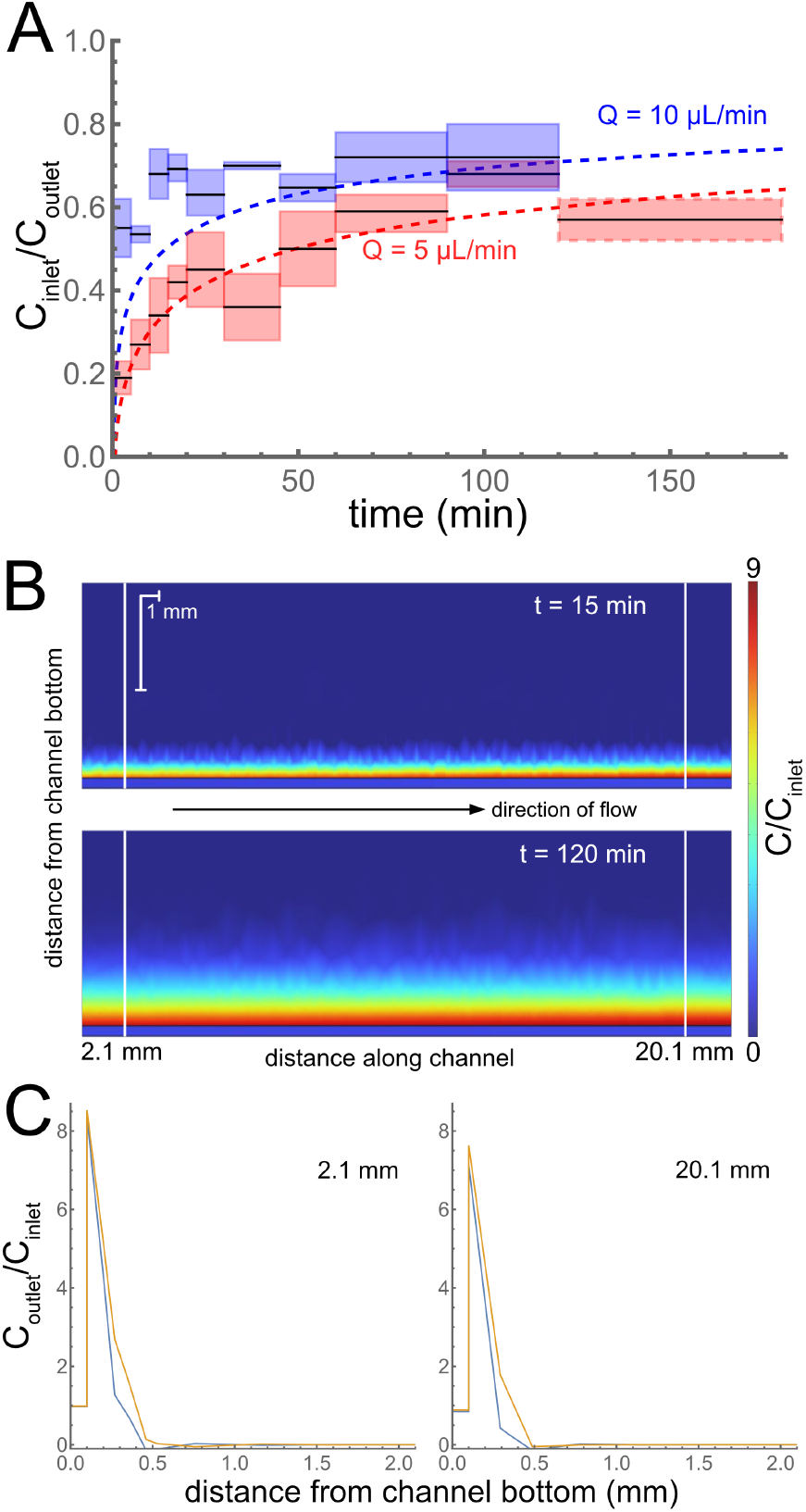
(A) Continuous flow experiments and simulations for indole solutions in microfluidic channels. Dashed lines represent COMSOL simulations. Boxes represent experimental data collected over the horizontal time span, with the box heights representing *±* one standard deviation. (B) 2D concentration profiles at 15 and 120 min after flow starts in a simulation of an indole solution flowing through a microfluidic channel at 5 µL/min. Horizontal and vertical length scales are different and follow from the 1-mm scale bars. (C) 1D concentration profiles taken at positions 2.1 and 20.1 cm along the channel (marked by vertical white lines in B).

## Discussion

We have investigated the PDMS interactions of ten chemicals, all reasonably hydrophobic (logP ≥ 1.9), but otherwise varying in their structure and properties. For the seven tested chemicals that interact with PDMS, we have fully characterized their interactions in a manner appropriate for use in 3D modeling, and confirmed that such modeling accurately predicts time-dependent doses in a PDMS-based microfluidic device. Within this small set of chemicals, the two key interaction parameters – the PDMS-water partition coefficient, K, and the diffusion constant in PDMS, D_*P*_ – each vary over several orders of magnitude. A mapping of these results into D_*P*_ -K parameter space is shown in Fig. 6A.

**Figure 6:**
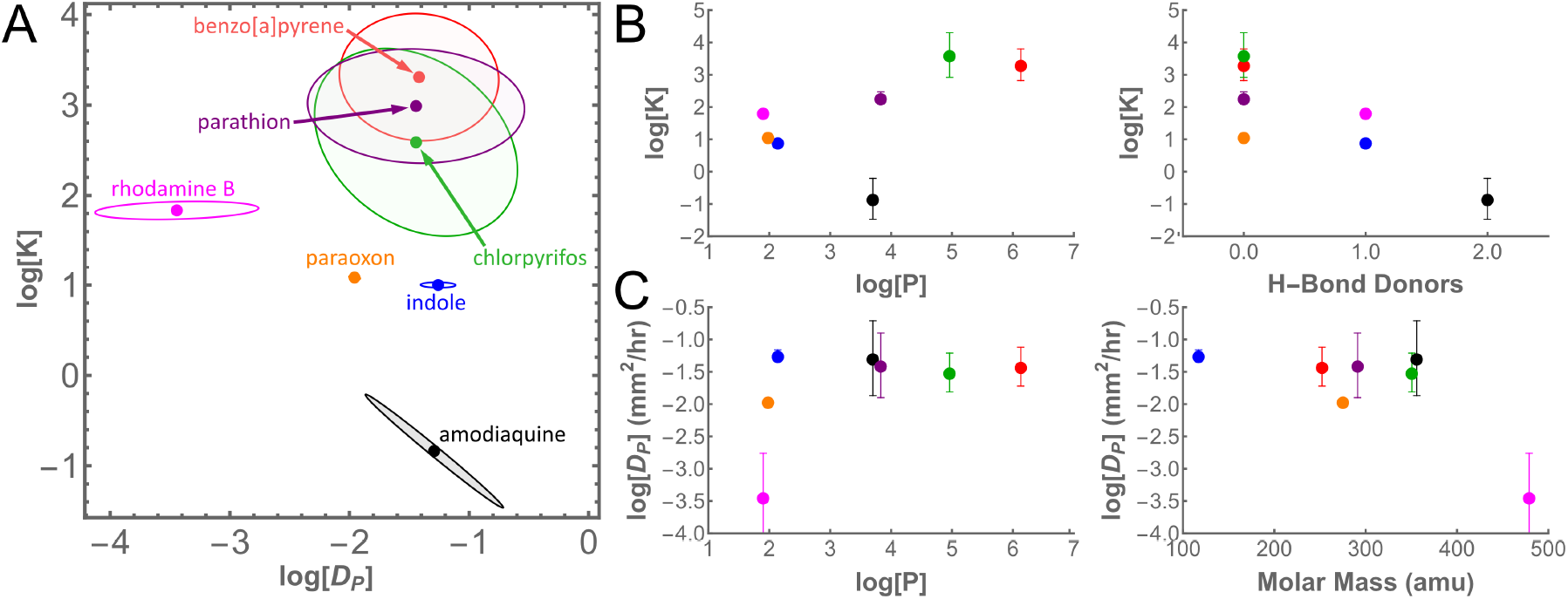
Correlations of PDMS-interaction parameters with select chemical properties. (A) Mapping of best-fit parameters for partitioning between water and PDMS (K) and diffusion in PDMS (D_*P*_). Note that results for benzo[a]pyrene, chlorpyrifos, and parathion were found via extrapolation from mixed solvents. Shaded areas represent 95% confidence limit ellipses. (B) Best-fit values of log K generally increase with hydrophobicity, as measured by logP, and decrease with the number of hydrogen bond donors. (C) Best-fit values of log D_*P*_ generally fall in a narrow band except for the much lower value for rhodamine B, which is also at the end of the tested range for the chemical properties logP and molar mass.

The best-fit parameter values for K and D_*P*_ display some interesting trends with chemical properties. The log K values, which range from around -1 to over 3, generally increase with the chemicals’ octanol-water partition coefficient logP (Fig. 6B); however, the result for amodiaquine is a notable exception, partitioning into PDMS much more weakly than the general trend suggests. Interestingly, amodiaquine also has more hydrogen-bond donor sites than any of the other chemicals tested, and the number of hydrogen-bond donors has been previously noted as a factor that decreases chemical binding to PDMS.[13] When the values of log K determined here are plotted against the number of hydrogen-bond donors, we find a similar decreasing trend (Fig. 6B). The trends for diffusivity in PDMS are less clear because six of the seven chemicals have D_*P*_ values within the same order of magnitude (0.01-0.1 mm^2^/hr). Interestingly, this range matches that for self-diffusion of PDMS chains.[36] In our test set, only rhodamine B stands alone as a slow diffusing outlier (4 × 10^*−*4^ mm^2^/h), and we confirmed its slow diffusion by direct imaging. Notably, rhodamine B lies at the lower end of our tested range for logP and at the upper end of the tested range for molar mass (Fig. 6C). Although our data set is too small to further delineate predictive correlations between molecular properties and chemical-PDMS interaction parameters, the observed trends suggest this as a logical avenue to pursue in the future.

On the other hand, one has to be careful in using simple read-across methods to predict chemical-PDMS interactions based on analogs: chemicals with similar molecular properties can display markedly different interactions with PDMS. In our test set, we have three chemical pairs that are reasonably similar to one other: fluorescein and FITC; rhodamine B and rhodamine 6G; and parathion and paraoxon. A fourth pair, FITC and amodiaquine, are not structurally similar, but FITC has been used previously as an analog to estimate amodiaquine’s PDMS-interaction parameters.[25] In the case of fluorescein and FITC, the molecules share a similar structure, and neither show any interaction with PDMS. In the case of rhodamine B and rhodamine 6G, the molecules are members of the same dye family with almost identical masses and structures; however, rhodamine B partitions into and diffuses through PDMS, while rhodamine 6G only binds to the PDMS surface. In the case of paraoxon and parathion, the molecules only differ by a single atom replacement of sulfur for oxygen. Despite this similarity, paraoxon partitions into PDMS much more weakly than parathion; the two compounds have partition coefficients that differ by two orders of magnitude. Finally, in the case of amodiaquine and FITC, the two compounds have very different structures, but share similar molecular weights and logP values. Based of these similarities, the fluorescent dye FITC has been previously used as an analog for estimating the PDMS interactions of amodiaquine.[25] Here, we measured the interactions directly for both compounds: FITC shows no interaction with PDMS in either disk soak, membrane, or optical diffusion experiments, but amodiaquine does partition into and diffuse within PDMS. Read-across methods may become better as we learn more about the molecular structure properties that determine chemical-PDMS interaction parameters, but analogs based solely on logP and molar mass are insufficient.

These cautionary analog results point to the importance of directly measuring PDMS interactions; however, many chemicals of interest may be too poorly soluble in water to conduct PDMS-interaction experiments in buffered aqueous solutions. Here, we show that one can instead take measurements in mixed solvents with high DMSO fractions and use a log-linear relationship to estimate the relevant parameters in water or in solutions between 1 and 5 percent DMSO by volume, a regime more compatible with the biological requirements of cells. Within our test set, this approach was done out of necessity for three compounds: benzo[a]pyrene, chlorpyrifos, and parathion. Both benzo[a]pyrene and chlorpyrifos are extremely hydrophobic and poorly soluble in water (solubilities of 1.62 µg/L and 1.4 mg/L, respectively, at 25 °C). The combination of poor aqueous solubility and low molar absorptivity (in the UV-Vis spectral region) required the addition of cosolvent to make disk-soak and membrane experiments feasible. For parathion, the necessity arose for a different reason. Parathion is more soluble in water (20 mg/L at 25 °C), and thus its concentration could be measured in aqueous media using UV-Vis spectroscopy; however, the partitioning of parathion into PDMS was so strong, that in experiments without a cosolvent, or with low fractions of DMSO (10% and 20% by volume), parathion reached its saturating concentration in PDMS (0.93 ng/mm^3^; Fig. S3). Since such saturation prevents measurement of a true partition coefficient, we needed experiments on parathion at higher cosolvent fractions than at first appeared necessary. We also conducted high-cosolvent experiments on indole, which is highly water soluble, but we did so only to explore the viability and validity of extrapolating partition coefficients with the log-linear relationship of equation (6). Based on the indole results, this extrapolation from high-cosolvent experiments appears valid and useful over the regimes investigated here.

## Conclusions

We have developed methods to characterize the chemical-PDMS interaction parameters needed to model the distribution of a chemical of interest when an aqueous solutions is in contact with PDMS. We have measured these physical parameters for ten hydrophobic chemicals, and confirmed the validity of these measurements through both direct observation of diffusion in PDMS and accurate predictions of time-dependent chemical loss in a PDMS-based microfluidic device. We find that partition coefficients and diffusion constants in PDMS can vary by orders of magnitude, and although there are trends with logP, with the number of hydrogen-bond donors, and with molar mass, one must exercise caution when using chemical analogs to infer PDMS-interaction parameters. With the battery of experiments and models presented here, one can measure chemical-PDMS interactions, even for highly hydrophobic compounds, and predictively model the time-dependent spatial distribution of a chemical of interest throughout the channels and PDMS bulk of a microfluidic device.

## Abbreviations

DMSO: dimethyl sulfoxide
FEM: finite element model(ing)
FITC: fluorescein-5-isothiocyanate
IVIVE: *in vivo*-*in vitro* extrapolation
PBS: phosphate buffered saline
PDMS: polydimethylsiloxane
OoC: organ-on-chip

## Definitions

c_*S*_ – concentration in solution

c_*S*_1 – concentration in source

c_*S*_2 – concentration in sink

c_*P*_ – concentration in PDMS

D_*S*_ – diffusivity in solution

D_*P*_ – diffusivity in PDMS

H – mass-transfer coefficient

K – partition coefficient

K_*P W*_ – PDMS-water partition coefficient

K_*P C*_ – PDMS-cosolvent partition coefficient

K_*P M*_ – PDMS-mixed solution partition coefficient

K_*P D*_ – PDMS-DMSO partition coefficient

f – volume fraction in solution

σ – Gaussian square width

## Data availability

Data for this article, including the results of disk and membrane soak experiments, and simulated solutions used in generating interpolation functions, are available at Open Science Framework at https://osf.io/892mf/.

## Acknowledgements

This publication was supported by U.S. EPA STAR Center Grant #84003101. Its contents are solely the responsibility of the grantee and do not necessarily represent the official views of the U.S. EPA. Further, U.S. EPA does not endorse the purchase of any commercial products or services mentioned in the publication. The authors would also like to thank Phillip Fryman for his technical assistance.

