## Supplemental Figures for "Toxicokinetics for organ-on-chip devices"

Nathaniel G. Hermann,[a] Richard A. Ficek,[a] Dmitry A. Markov, [b] Lisa J. McCawley,[b] and M. Shane Hutson[a]*
[a] *Department of Physics and Astronomy, Vanderbilt University*
[b] *Department of Biomedical Engineering, Vanderbilt University*

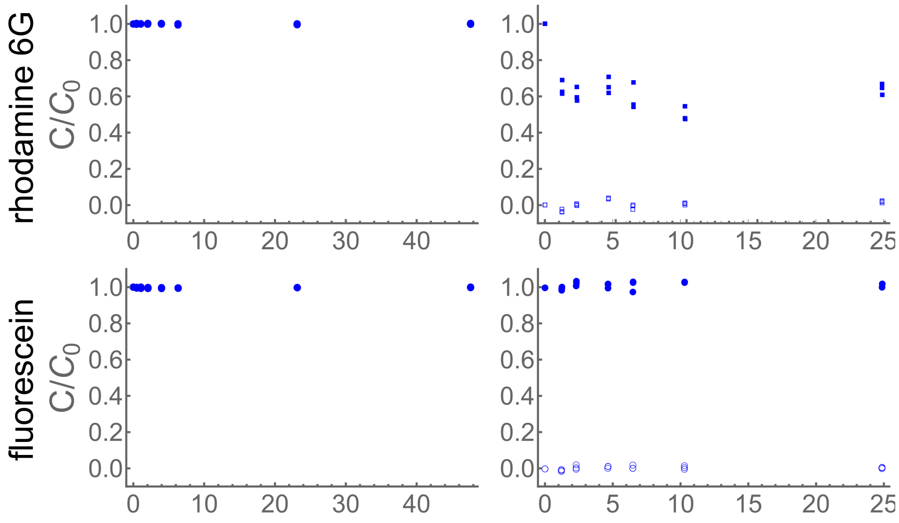


Figure S1: Disk-soak and diffusion-through-membrane results for two additional chemicals in phosphate-buffered saline: rhodamine 6G binds to the surface of PDMS, but does not diffuse into the bulk; fluorescein does not measurably interact with PDMS. Left column shows disk-soak data and associated fits (dashed). Right column shows diffusion-through-membrane results with data from both source chamber (filled symbols) and sink chamber (open symbols) with associated fits (dashed).

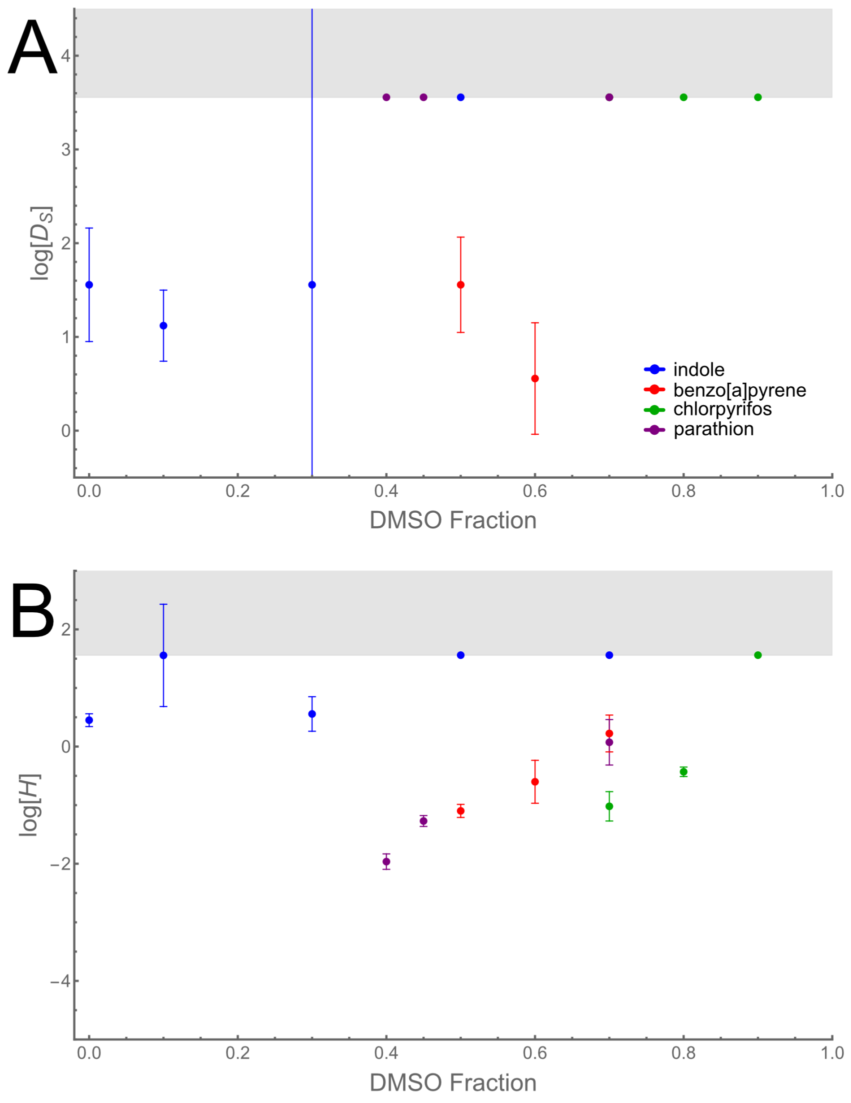


Figure S2: Best-fit values for the diffusion constants in solution, $\log D_S$ (A), and the mass-transport coefficient, $\log H$ (B), for indole, benzo[a]pyrene, chlorpyrifos, and parathion at several DMSO volume fractions. Shaded regions mark regimes in which the plotted parameter is not rate limiting. Note that in (A), the rate-limiting boundary ($\log D_S \approx 3.55$) has several overlapping points for different chemicals.

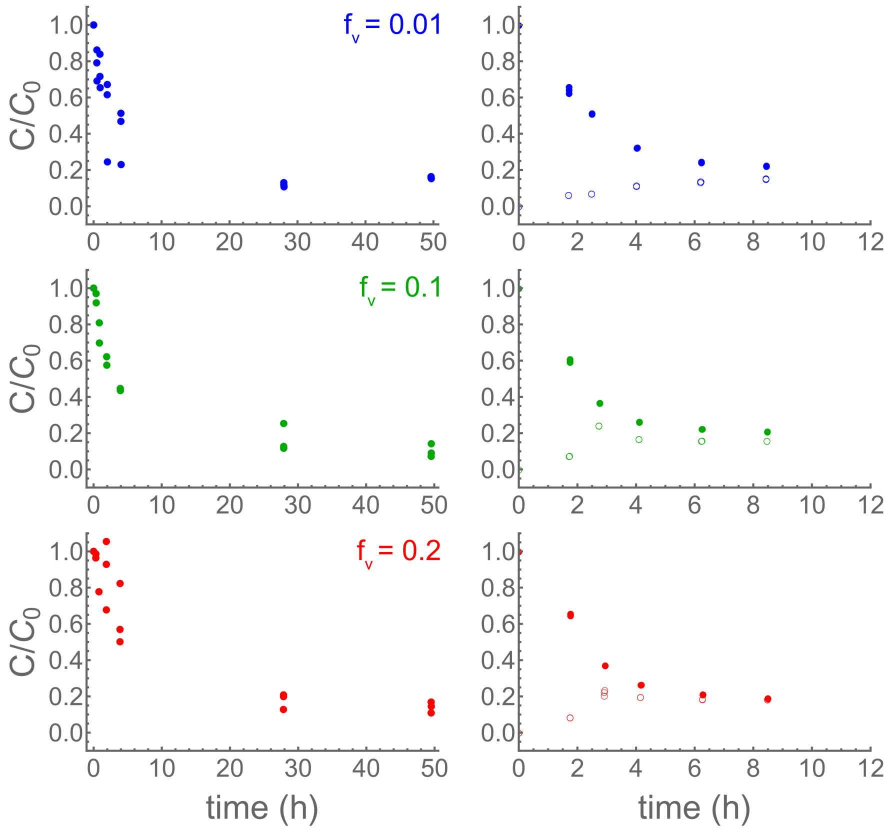


Figure S3: Experimental results from disk-soak (left) and diffusion-through-membrane experiments (right) for parathion in mixed PBS-DMSO solutions at the noted DMSO volume fractions, $f_V$. For disk soaks, the equilibrium concentration in all three mixed solutions is around 10% of the initial concentration; for membrane experiments, the source and sink chambers equilibrate around 20% of the initial concentration. These results indicate a volume saturation of parathion in PDMS at an approximate concentration of 3.2 mM, or 0.93 ng/mm$^3$.

### Supplemental Movies

Figure S4: (Left) Direct observation of diffusion of rhodamine B into bulk PDMS. A microchannel was pre-soaked with a rhodamine B solution for three hours to pre-load chemical into the PDMS walls. The microchannel was then flushed, and the subsequent diffusion of pre-loaded dye was imaged for 12 hours. (Right) Under similar conditions, rhodamine 6G remains on the PDMS surface and does not measurably diffuse into bulk PDMS over 12 hours.

Figure S5: Sample results from a COMSOL Multiphysics simulation of the partitioning and diffusion of indole from a solution flowing through a microfluidic channel. Flow rate was 5 $\mu$L/min over a duration of two hours, with flow direction from left to right along the bottom of the figure. Highest concentrations of indole are red; lowest are blue.
